# Ca^2+^-phospholipid–dependent regulation of Munc13-1 is essential for post-tetanic potentiation at mossy fiber synapses and supports working memory

**DOI:** 10.1101/2025.08.03.668318

**Authors:** Francisco José López-Murcia, Dilja Krueger-Burg, Sally Wenger, Tania López-Hernández, Noa Lipstein, Holger Taschenberger, Nils Brose

## Abstract

Hippocampal mossy fiber (hMF) to CA3 pyramidal cell synapses are thought to support the formation of working memory through presynaptic short-term facilitation (STF) and post-tetanic potentiation (PTP). However, the molecular mechanisms underlying these transient forms of synaptic enhancement remain poorly understood. We show here that Munc13-1-mediated priming of synaptic vesicles (SVs) at active zones controls hMF STF and PTP in response to Ca^2+^-phospholipid and Ca^2+^-calmodulin (CaM) signaling. Knock-in mice expressing Munc13-1 variants that are insensitive to Ca^2+^-phospholipid and Ca^2+^-CaM signaling exhibit severely impaired STF and PTP at hMF synapses. Moreover, the PTP-induction threshold is strongly increased upon the loss of Ca^2+^-phospholipid-Munc13-1 signaling. Since these synaptic defects are accompanied by working memory deficits, especially in mice expressing the Ca^2+^-phospholipid-insensitive Munc13-1 variant, we conclude that Ca^2+^-dependent regulation of Munc13-1-mediated SV priming co-determines hMF short-term plasticity and working memory formation.

## INTRODUCTION

Hippocampal mossy fiber (hMF) → CA3 pyramidal cell synapses, key information processing units in the hippocampal network^1^, exhibit pronounced synaptic short-term facilitation (STF) and post-tetanic potentiation (PTP) upon repetitive presynaptic action potential (AP) firing. These types of short-term plasticity (STP) are thought to play crucial roles in working memory formation and other brain functions^2–4^ but their molecular underpinnings have yet to be determined.

Presynaptic Ca^2+^-buffer saturation^5–7^ and certain paralogs of the Synaptotagmin family of Ca^2+^-sensor proteins^8^ contribute to STF in hMF synapses. On the other hand, PTP is associated with an increased number of docked SVs at active zones (AZs) in hMF^9^ and other brain synapses^10,11^. While several Ca^2+^-dependent kinases were implicated in inducing PTP^12–14^ and frequency facilitation (FF)^15^, the proteins that ultimately execute FF and PTP are unknown. Components of the SV priming machinery are prime candidates because the Ca^2+^-dependent acceleration of SV priming is known to regulate STP^16–19^.

Munc13-1 is an essential SV priming protein^20–22^. Its activity is regulated by three distinct pathways – Ca^2+^-calmodulin (CaM) signaling via its CaM-binding domain^23,24^, Ca^2+^-phospholipid signaling via its C_2_B domain^16,25^, and diacylglycerol (DAG) signaling via its C_1_ domain^26,27^. In cultured autaptic neurons^23,25,27^ and at calyx of Held synapses^16,17^, the regulation of Munc13-1 activity via these signaling pathways modulates short-term depression (STD).

In the present study, we tested whether Munc13-1-mediated SV priming and its Ca^2+^-dependent regulation participates in synaptic short-term enhancement processes, such as STF and PTP, and whether these mechanisms contribute to working memory function. Using knock-in (KI) mutant mice expressing Munc13-1 variants that are insensitive to Ca^2+^-CaM signaling^17^ or Ca^2+^-phospholipid signaling^16^, we show that inactivation of these signaling pathways impairs paired-pulse facilitation (PPF), FF, and, strikingly, also PTP in hMF → CA3 pyramidal cell synapses. We further demonstrate that the Ca^2+^-dependent enhancement of Munc13-1-mediated SV priming supports working memory formation.

## RESULTS

### Activation of Munc13-1 C_2_B and Ca^2+^-CaM binding domains is required for STF in hMF synapses

To assess whether the C_2_B and the CaM-binding domains of Munc13-1 participate in the activity-dependent regulation of hMF plasticity, we used two KI mouse lines, (i) the DN line, which expresses a Munc13-1^D705,711N^ variant that lacks Ca^2+^-dependent phospholipid binding to the C_2_B domain^16,25^, and (ii) the WR line, which expresses a Munc13-1^W464R^ variant that lacks Ca^2+^-CaM binding^17^ (**Figure 1A**). Immunofluorescence microscopy of hippocampal sections revealed wild-type (wt) like Munc13-1 abundance and distribution in hMF terminals of both KI mouse lines (**Figure S1D**)^16,17^.

**Figure 1.**
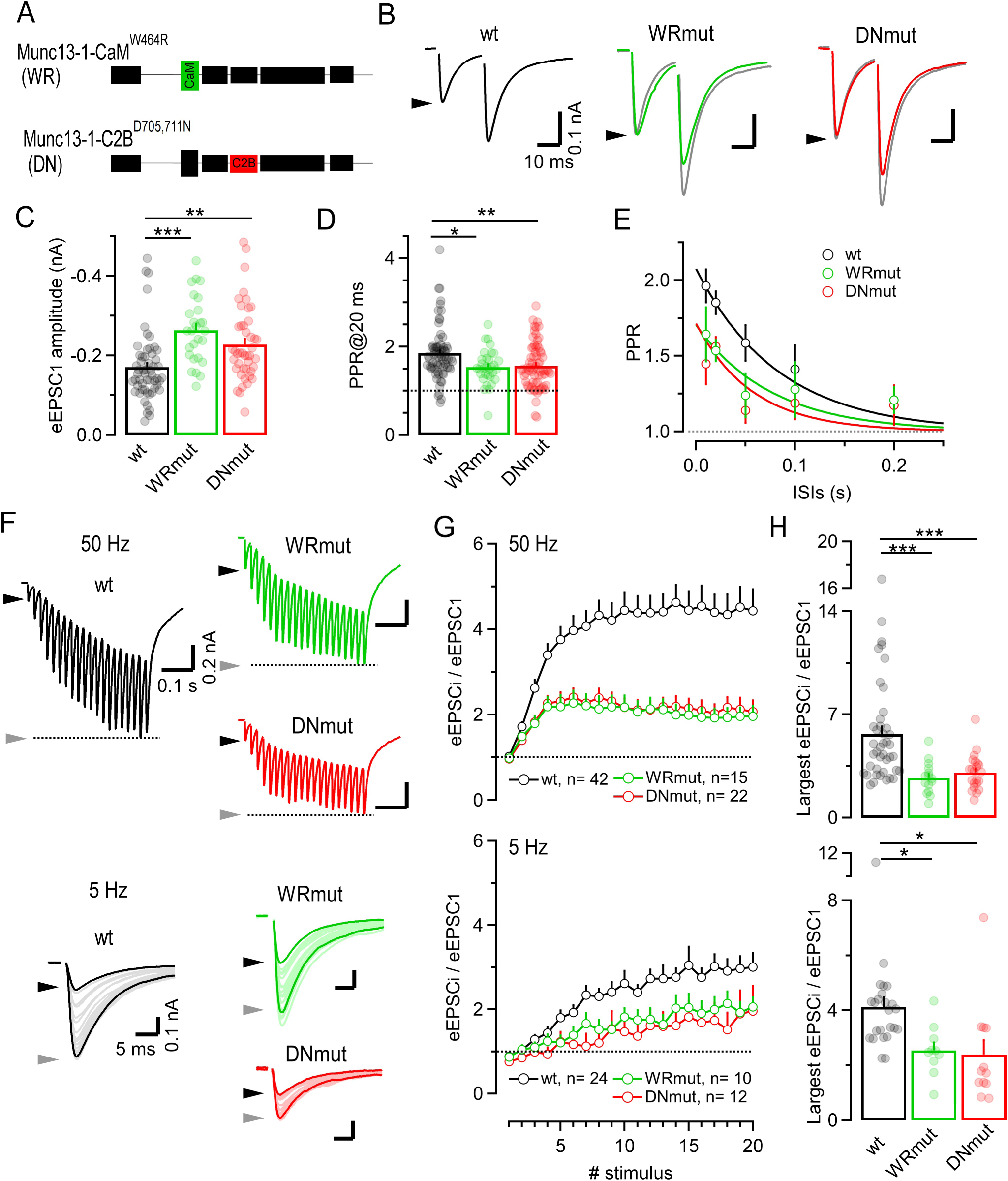
Paired-pulse and frequency facilitation are impaired in WRmut and DNmut hMF synapses. **(A)** Schematic of wt and mutant Munc13-1 protein domains. **(B)** Average putative unitary hMF eEPSC pairs (ISI = 20 ms) recorded in CA3 pyramidal neurons of wt (n=17, *black*), WRmut (n=10, *green*), and DNmut (n=21, *red*) mice. Arrowheads indicate mean eEPSC_1_ amplitudes. For comparison, peak scaled versions of the average wt trace (*grey traces*) are superimposed onto WRmut and DNmut eEPSCs. **(C-D)** Bar graphs and scatter dot plots displaying means and individual values, respectively, of eEPSC_1_ amplitudes (C; average over all trains recorded in each synapse) and PPRs (D; ISI = 20 ms). wt, n=56 and 68; WRmut, n= 27 and 32; and DNmut, n=44 and 69. Dotted line indicates unity. **(E)** PPRs plotted against ISI for wt (*black*), WRmut (*green*), and DNmut (*red*) hMF synapses. Data were fitted with mono-exponential curves (*solid lines*; τ_wt_ = 83.4 ms, τ_WRmut_ = 76.4 ms, and τ_DNmut_ = 57.4 ms). Dotted line indicates unity. **(F)** Average hMF eEPSCs trains elicited by 50 Hz (*top*) and 5 Hz (*bottom*) stimulation recorded in wt (*black*; n = 37 and n = 23), WRmut (*green*; n = 14 and n = 10), and DNmut (*red*; n = 24 and n=12) synapses, respectively. Black arrowheads mark eEPSC_1_ amplitudes. Gray arrowheads mark either the largest eEPSC peak amplitudes (50 Hz) or eEPSC_20_ amplitudes (5 Hz). **(G)** Normalized amplitudes of 50 Hz (*top*) and 5 Hz (*bottom*) eEPSC trains. For each synapse, eEPSC amplitudes were normalized to the average eEPSC_1_ amplitude of all eEPSC trains. Dotted lines indicate unity. **(H)** Bar graphs and scatter dot plots showing means and individual values, respectively, of the largest normalized eEPSC amplitudes during 50 Hz (*top*) and 5 Hz (*bottom*) stimulation. Error bars indicate SEM. (ANOVA followed by Tukey-test, * p<0.05, ** p<0.01, *** p<0.001).

We recorded putative unitary excitatory postsynaptic currents (eEPSCs) at near-physiological temperature in CA3 pyramidal neurons. Pharmacologically isolated eEPSCs were elicited by focal extracellular stimulation of mossy fiber axons (**Figure S1A-S1C**). hMF synapses of DN and WR wt littermate mice were functionally indistinguishable (**Figures S1E-S1G)**, and therefore the corresponding data were pooled.

Initial eEPSCs were slightly but consistently larger in WRmut and DNmut synapses as compared to wt (**Figures 1B and 1C**)^16,17,28^. Paired-pulse ratios (PPR = EPSC_2_/EPSC_1_; using 0.01–0.2 s inter-stimulus intervals [ISIs]), were significantly lower in WRmut and DNmut synapses in comparison to wt synapses (**Figures 1D and 1E**), indicating diminished paired-pulse facilitation (PPF).

We next examined FF, a hallmark of short-term plasticity at hMF synapses, during short trains (20 stimuli, 1–50 Hz) (**Figure S1H**). For all frequencies tested, the magnitude of FF, quantified as the ratio of the largest over the initial eEPSC, was significantly reduced in WRmut and DNmut synapses in comparison to wt synapses (e.g. at 5 Hz and 50 Hz: wt, ∼4-fold and ∼5.7-fold; WRmut, ∼2.5-fold and ∼2.7-fold; DNmut, ∼2.4-fold and ∼3-fold; **Figures 1F-1H**). FF was almost undetectable in WRmut and DNmut synapses at the lowest frequency (1 Hz; **Figure S1H**). In addition, the buildup of synaptic enhancement during stimulus trains^29^ at frequencies between 1–20 Hz occurred more slowly in mutant hMF synapses, especially in DNmut mice (**Figure S1I**).

In sum, these experiments demonstrate that Ca^2+^-phospholipid and Ca^2+^-CaM regulation of Munc13-1 co-determine magnitude and time course of STF in hMF synapses.

### Facilitation of hMF synapses by low-frequency conditioning requires Ca^2+^-phospholipid and Ca^2+^-CaM signaling to Munc13-1

Dentate gyrus granule cells fire at low baseline rates but intermittently discharge brief high-frequency bursts. At low baseline firing rates, hMF → CA3 pyramidal cell synapses typically fail to evoke postsynaptic APs, while burst firing profoundly increases synaptic strength^9^. To test whether low-frequency conditioning transiently facilitates hMF → CA3 transmission and whether Munc13-1-mediated SV priming contributes to this effect, we recorded 100 Hz eEPSC trains before (ctrl) and after (cond) conditioning by low-frequency stimulation (LFS; 20 stimuli at 2 Hz). Such brief LFS is expected to induced only minimal summation of AP-evoked presynaptic Ca^2+^ transients (τ ≍ 490 ms^30,31^), thereby limiting the contribution of Ca^2+^-buffer saturation to synaptic facilitation. 100 Hz trains were limited to 15 stimuli to minimize confounding effects of polysynaptic “recurrent” excitation of neighboring CA3 pyramidal neurons^32^. Pre-conditioned 100 Hz trains were recorded either 0.5 s or 5 s following LFS (Δt= 0.5 s or 5 s; **Figures 2A**).

**Figure 2.**
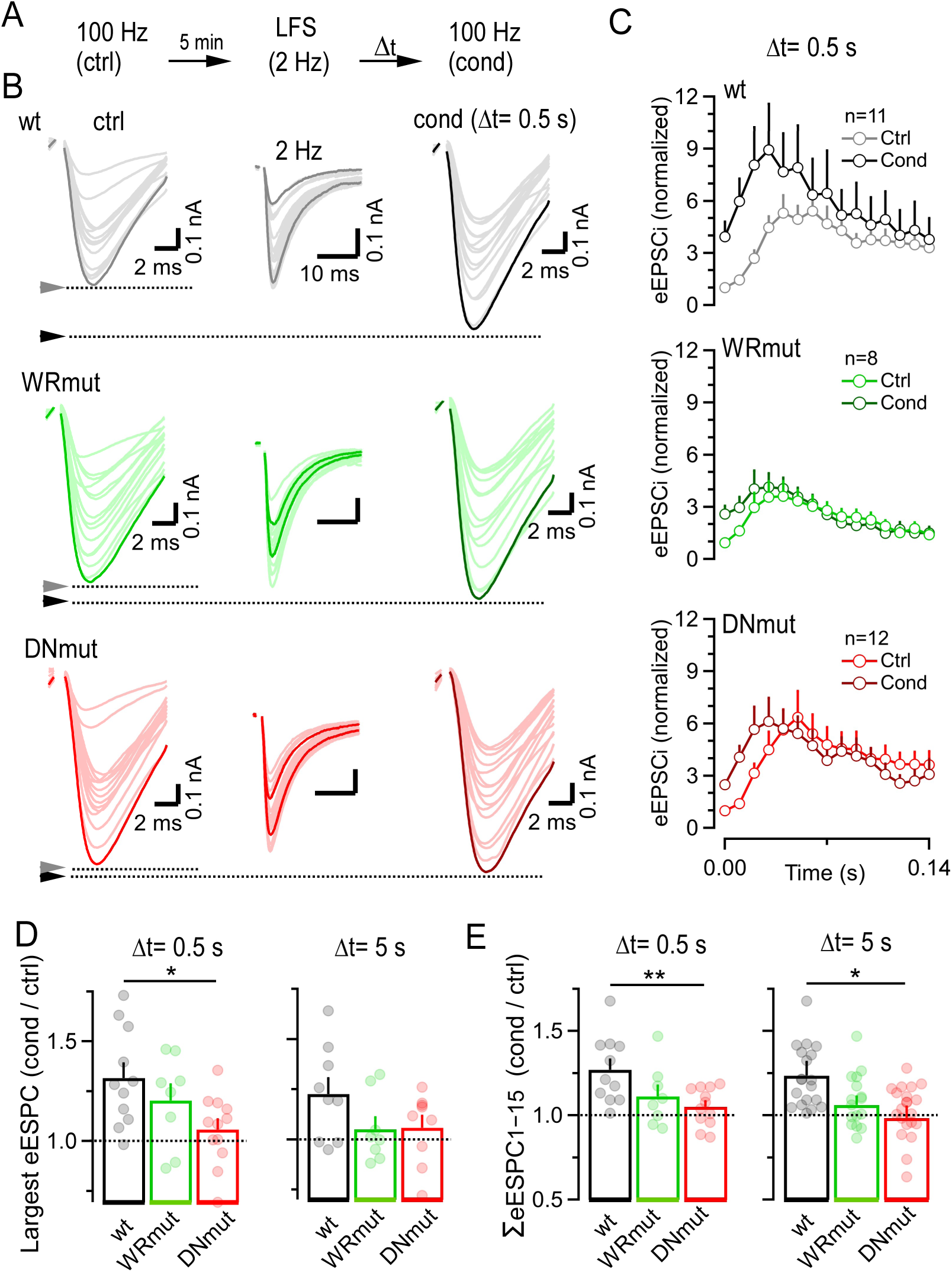
Synaptic facilitation induced by low-frequency conditioning is impaired in WRmut and DNmut hMF synapses. (**A**) Experimental paradigm: 15 stimuli at 100 Hz were applied (control eEPSC train, ctrl, followed by a five-minute recovery period, after which synapses were conditioned using low-frequency stimulation (LFS; 2 Hz, 20 stimuli) to induce facilitation. Shortly after (Δt = 0.5 s or 5 s) conditioning, a second 100 Hz train was recorded (conditioned eEPSC train, cond). This stimulation protocol was repeated up to three times at five-minute intervals. (**B**) Average amplitudes (eEPSC_1-15_) of ctrl (*left column*) and cond (*right column*) 100 Hz eEPSC trains recorded in wt (*black*, n = 11), WRmut (*green*, n = 8) and DNmut (*red*, n = 12) hMF synapses at Δt = 0.5 s. The largest mean eEPSCs in the trains are highlighted (*bold traces*) and their peak amplitudes are indicated by arrowheads. eEPSCs of the LFS train are shown for comparison (*middle column*) with eEPSC_1_ and eEPSC_20_ highlighted (*bold traces*). (**C**) Mean normalized eEPSC_1-15_ amplitudes of ctrl eEPSC trains (*light colors*) and cond eEPSC trains (*solid color*) in wt (*black*), WRmut (*green*), and DNmut (*red*) hMF synapses Δt = 5 s. eEPSC amplitudes were normalized to the average eEPSC_1_ amplitude of the corresponding ctrl 100 Hz and LFS 2 Hz trains. **(D-E)** Bar graphs and scatter dot plots showing averages and individual values, respectively, of the ratios of the largest eEPSC amplitudes of the cond trains over that of the respective values of the ctrl trains (D) and of the cumulative amplitudes of cond trains and the corresponding values of the ctrl trains (E), for Δt = 0.5 s (*left*) or Δt = 5 s (*right*). Dotted lines indicate the unity. Error bars represent SEM. (ANOVA followed by Tukey-test, *p<0.05, **p<0.01)

During LFS, wt synapses facilitated ∼3-fold. Consequently, the initial eEPSC amplitudes of cond trains were larger than those of the corresponding ctrl trains, and eEPSCs further facilitated during the 100 Hz train. In wt hMF synapses, the maximum eEPSC recorded during the cond train was on average 9-fold larger than the initial eEPSC of the ctrl train (eEPSC_1,ctrl_; **Figure 2C**). When comparing the largest eEPSCs during cond and ctrl trains, we observed on average a ∼30% increase at Δt= 0.5 s (683 ± 55 pA [ctrl] vs. 964 ± 105 pA [cond]) and a ∼24% increase at Δt= 5 s (946 ± 143 pA [ctrl] vs. 1174 ± 227 pA [cond]) (**Figures 2B and 2D and S2A**). LFS-induced facilitation of hMF → CA3 transmission was also evident when analyzing the ratios of the cumulative eEPSC_1-15_ amplitudes of the 100 Hz trains recorded before and after pre-conditioning (**Figure 2E and S2C**). LFS-induced facilitation gradually decayed with longer recovery intervals with a time constant of ∼20–30 s (**Figures S2A-S2C**).

To assess whether Munc13-1 contributes to LFS-induced facilitation, we examined the extent of facilitation relative to eEPSC_1,ctrl_ and the ratios of the largest eEPSC amplitudes during ctrl and cond 100 Hz trains in Munc13-1 KI mice at Δt= 0.5 s and 5 s (**Figure 2C and 2D**). At Δt= 0.5 s, LFS-induced facilitation was significantly reduced in WRmut synapses (∼14%; 898 ± 229 [ctrl] vs. 1047 ± 248 pA [cond]) and abolished in DNmut synapses (1026 ± 127 pA [ctrl] vs. 1029 ± 149 pA [cond]; **Figures 2D, S2D and S2E**). At Δt = 5 s, neither mutant showed LFS-induced facilitation. Analysis of cumulative eEPSC_1-15_ amplitudes corroborated these results (**Figures 2E**).

Taken together, these observations indicate that LFS-induced facilitation of hMF → CA3 transmission is a Munc13-1-dependent form of STF, requiring Ca^2+^-phospholipid and Ca^2+^-CaM signaling.

### Ca^2+^-phospholipid and Ca^2+^-CaM signaling to Munc13-1 are required for PTP

Post-tetanic potentiation is a prominent form of presynaptic enhancement of glutamate release at hMF → CA3 pyramidal cell synapses, which exhibit a remarkably low PTP-induction threshold^33^. Since overfilling of the pool of fusion-competent SVs was recently proposed to critically contribute to PTP^9^, we next tested whether Ca^2+^-phospholipid and Ca^2+^-CaM signaling to Munc13-1 are involved in PTP induction.

PTP was induced by high-frequency stimulation (HFS; 50 Hz) consisting of 5, 10, 20, 50, or 70 stimuli. This stimulation frequency resembles the physiological firing rates of granule cells during AP bursts^9,34,35^. Changes in glutamate release were assessed by recording eEPSC pairs (eEPSC_1_ and eEPSC_2_; 20 ms ISI; PP@20ms) every 20 s before and after HFS. The mean of all eEPSC_1_ amplitudes recorded before HFS was defined as the baseline eEPSC amplitude (eEPSC_basal_). The PTP magnitude was defined as the ratio of the amplitudes of eEPSC_1_ recorded immediately after HFS over that of eEPSC_basal_ (**Figure 3A**).

**Figure 3.**
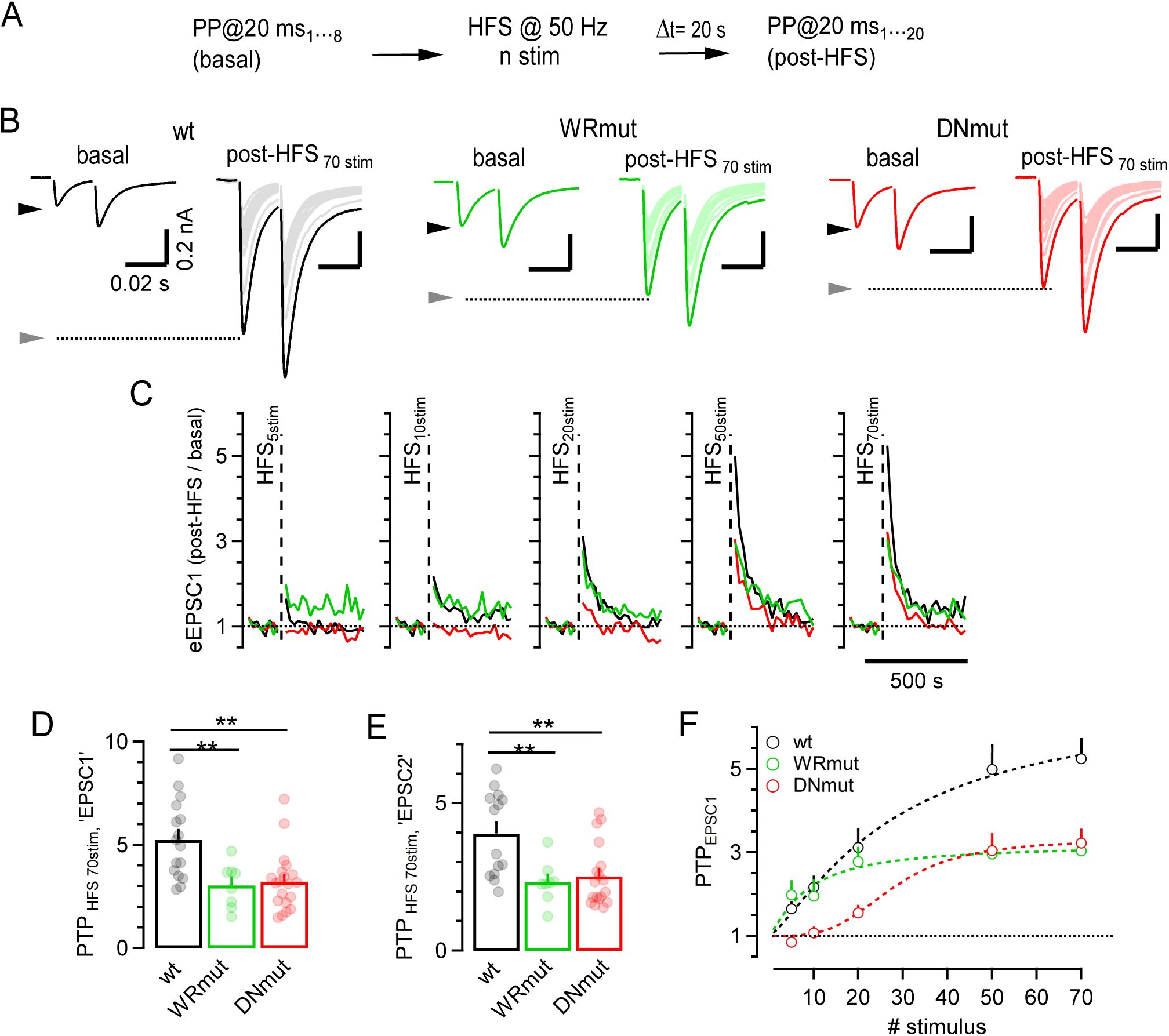
PTP magnitude is reduced in WRmut and DNmut hMF synapses. (**A**) Experimental paradigm: eEPSCs pairs (ISI = 20 ms) were recorded every 20 s before and after PTP induction in wt, WRmut, and DNmut hMF synapses. The basal eEPSC_1_ amplitude represents the average of eight eEPSCs pairs recorded before PTP induction. High-frequency stimulation (HFS) PTP induction trains (50 Hz) consisted of 5–70 stimuli (HFS_5stim_–HFS_70stim_). Following HFS, another twenty eEPSC pairs (ISI = 20 ms) were recorded at 20 s intervals. The ratio of the first eEPSC_1_ amplitude measured after HFS (Δt = 20 s) over the corresponding basal eEPSC_1_ amplitude represents the PTP magnitude. The same analysis was applied to eEPSC_2_. (**B**) Average basal eEPSC pairs recorded before (*left*) and average potentiated eEPSC pairs recorded at various times after HFS_70stim_ in wt (*black*, n = 16), WRmut (*green*, n = 8) and DNmut (*red*, n = 19) hMF synapses. The first eEPSC pair recorded after HFS is emphasized (*bold traces*). Arrowheads and dashed lines indicate peak eEPSC amplitudes. (**C**) Time course of normalized eEPSC_1_ amplitudesbefore and after HFS (*dashed lines*). Error bars are omitted for clarity. Dotted and dashed lines represents unity and stimulation onset, respectively. **(D-E)** Bar graphs and scatter dot plots showing mean and individual values, respectively, of PTP magnitudes for eEPSC_1_ (*left*) and eEPSC_2_ (*right*) in wt (*black*), WRmut (*green*) and DNmut (*red*) hMF synapses. **(F)** PTP magnitudes for eEPSC_1_ plotted as a function of the number of stimuli applied during HFS. Experimental data were fitted with a Michaelis-Menten-like dose-response relationship (wt and WRmut) or with a Hill equation (DNmut; Hill coefficient (n) = 3.5). Dashed and dotted lines represents fit results and unity, respectively. Error bars represent SEM (ANOVA followed by Tukey-test, ** p<0.01).

In wt synapses, five stimuli were sufficient to induce substantial potentiation (**Figures 3C and S3A)**, whereas DNmut synapses required at least 20 stimuli. Moreover, the PTP magnitude was reduced for all induction train durations in DNmut synapses (**Figure S3A)**. WRmut synapses, in contrast, showed wt-like PTP magnitude for short induction trains (≤10 stimuli) but reduced PTP for longer ones (≥20 stimuli) (**Figures 3B, 3C and S3A**). In wt and mutant synapses, PPRs measured immediately after HFS were smaller than those measured prior to PTP induction, consistent with previous studies^29^. Reduced PTP was observed in mutant synapses regardless of whether EPSC_1_ or EPSC_2_ was analyzed to quantify PTP (**Figures 3D, 3E**).

The relationship between PTP magnitude and number of stimuli in the induction train was well described by a Michaelis-Menten like function in wt and WRmut synapses. Impaired Ca^2+^-CaM regulation of Munc13-1 primarily diminishes the PTP magnitude. In contrast, a sigmoid function was required to described the dose-response relationship in DNmut synapses (**Figure 3F**), reflecting the increased PTP-induction threshold in addition to the reduced PTP magnitude.

In summary, these findings demonstrate a crucial role of Ca^2+^-phospholipid and Ca^2+^-CaM signaling to Munc13-1 in PTP induction at hMF → CA3 pyramidal cell synapses.

### Abolished Ca^2+^-phospholipid signaling to Munc13-1 at hMF synapses interferes with spatial working memory formation

Working memory was long believed to be stored in the form of persistent neuronal activity, but this notion has been challenged in recent years by findings that synaptic alterations, including presynaptic forms of STP such as synaptic facilitation and PTP, may also play important roles^2,9,36–38^. However, support for this hypothesis originates primarily from computational models, while experimental evidence is scarce. To examine experimentally whether Ca^2+^-phospholipid and Ca^2+^-CaM signaling to Munc13-1 contribute to working memory, we assessed the performance of DN and WR KI mutant mice in a spatial working memory task, the 8-arm radial maze (8ARM).

Prior to the 8ARM task, mice underwent visual cliff (VC) and open field (OF) testing to rule out confounding deficits in vision, locomotion, or anxiety (**Figure S4A-S4C**). WRmut mice were similar to wt littermates across all tested parameters (**Figures S4D, S4F and S4H**). DNmut mice displayed normal visual and locomotor function, along with a mild reduction in anxiety-related behavior, which is, however, unlikely to impact 8ARM performance (**Figures S4E, S4G and S4I)**.

In the 8ARM task, mice were required to search for food pellets located in reward zones at the ends of three fixed arms (**Figure 4A**). Each of 10 daily sessions comprised four trials, during which mice were expected to collect all three pellets while avoiding revisiting the reward zones after pellet retrieval. Re-entries into previously visited reward zones were scored as working memory (WM) errors (**Figures 4B and 4C**).

**Figure 4.**
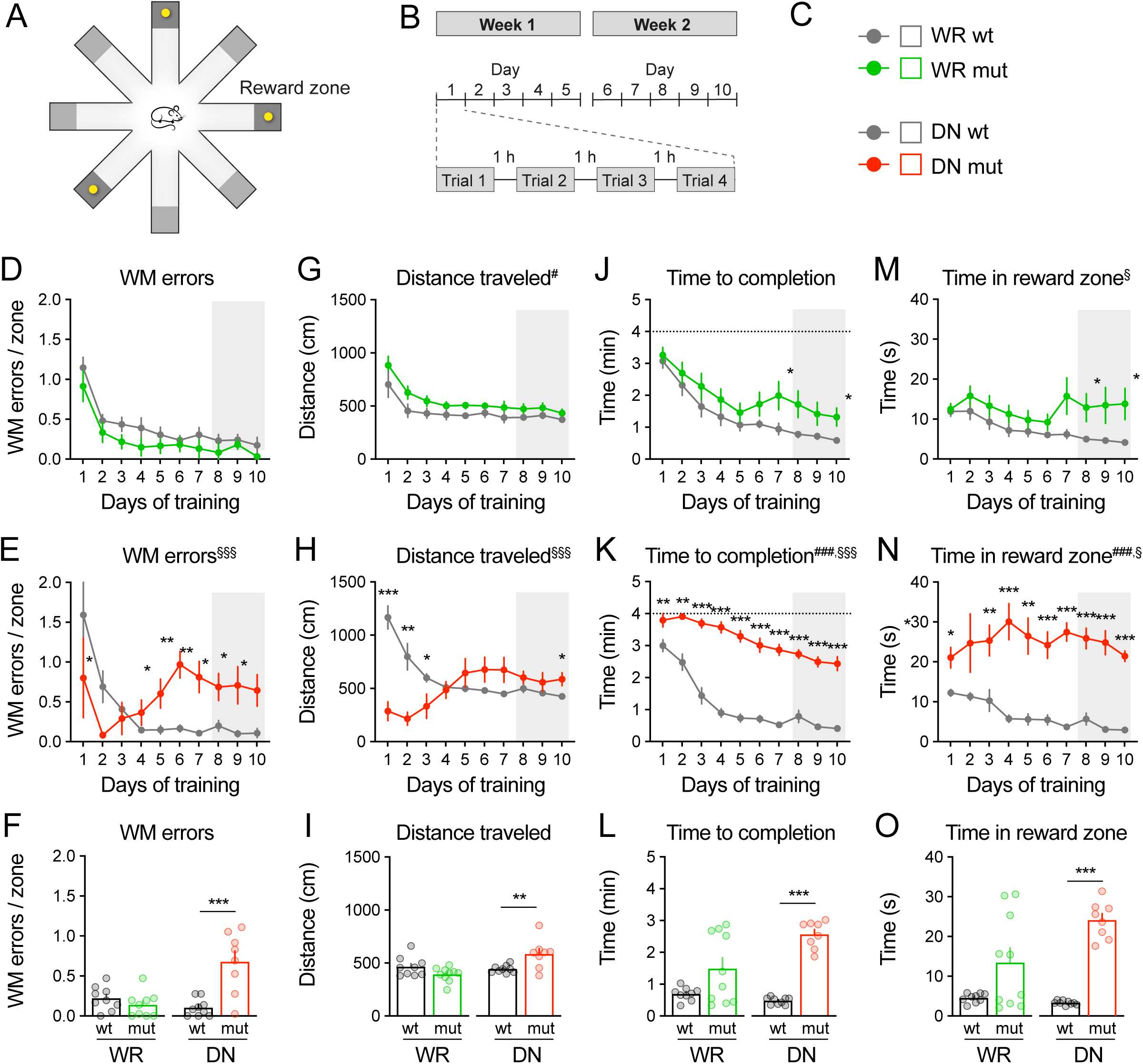
Mutations in Munc13-1 C_2_B and CaM-binding domains disrupt working memory. **(A)** Schematic representation of the 8-arm radial maze, indicating the location of the reward zones and a sample distribution pattern of the three food pellets (yellow circles). **(B)** Schematic representation of the behavioral paradigm, indicating the number of daily sessions per week and the number of trials per daily session. **(C)** Representation of the behavioral cohorts tested (all male mice, KI mice and their corresponding wt littermates, 8–12 weeks old at start of experiment). **(D-F)** Number of working memory errors per reward zone, defined as re-entries into the reward zone, for WR (D) and DN (E) mouse lines across all 10 daily sessions, and averages across the final 3 daily sessions (F). **(G-I)** Total distance travelled (in cm) in order to retrieve all three food rewards for WR (G) and DN (H) mouse lines across all 10 daily sessions, and averages across the final 3 daily sessions (I). **(J-L)** Total time (in min) taken to retrieve all three food rewards for WR (J) and DN (K) mouse lines across all 10 daily sessions, and averages across the final 3 daily sessions (L). **(M-O)** Total time (in min) taken to retrieve all three food rewards for WR (M) and DN (N) mouse lines across all 10 daily sessions, and averages across the final 3 daily sessions (O). D, E, G, H, J, K, M and N: Two-way ANOVA with repeated measures for day of training (main effect of day, not shown; main effect of genotype, ^#^ p<0.05, ^##^ p<0.01, ^###^ p<0.001; genotype x day interaction, ^§^ p<0.05, ^§§§^ p<0.001), followed by Tukey’s *posthoc* test (* p<0.05, ** p<0.01). F, I, L, and O: Unpaired two-tailed Student’s t-test or Mann-Whitney test; *p<0.05, **p<0.01, ***p<0.001). Error bars represent SEM.

DN and WR wt littermate mice learned to efficiently collect rewards with minimal WM errors during the final training sessions (**Figures 4D-4F**, *grey*). WRmut mice performed comparably to wt controls, with no significant differences in WM errors or total distance travelled (**Figures 4D, 4F, 4G and 4I**), but exhibited a modest increase in time spent in rewarded zones in later sessions, resulting in a slight delay in trial completion (**Figures 4J, 4L, 4M and 4P**). In contrast, DNmut mice displayed a striking behavioral phenotype. During the first three training days, these mice showed minimal exploration and rarely collected food, remaining largely immobile in one arm or food zone (**Figure 4H and 4K**), despite exhibiting normal locomotion and reduced anxiety in prior testing (**Figure S4E-S4K**). Accordingly, they committed few WM errors early on, since many food rewards were not retrieved at all. However, from day four onward, DNmut mice developed marked deficits in reward-related behavior. After retrieving a pellet, they spent significantly more time in the food zone, seemingly continuing to search for a reward that had already been consumed (**Figures 4K, 4L, 4N, and 4O**). Moreover, after eventually exiting the rewarded food zone, DNmut mice frequently returned to the same reward zone, resulting in a large number of WM memory errors (**Figures 4E and 4F**), This behavior indicates a pronounced inability to register or remember successful reward retrieval.

On aggregate, these findings indicate that Ca^2+^-dependent regulation of Munc13-1 is critical for the encoding or maintenance of recent reward-related information, supporting the recent notion that presynaptic forms of short-term plasticity play a key role in working memory processing.

## DISCUSSION

Activity-dependent upregulation of Munc13-1-mediated SV priming counteracts synaptic depression due to SV depletion during sustained presynaptic activity^16,17^. Here, we show that Munc13-1 also contributes to multiple forms of short-term facilitation, including PPF, FF, and PTP, at hMF → CA3 pyramidal cell synapses. Disrupting Ca^2+^-phospholipid or Ca^2+^-CaM signaling to Munc13-1 severely impairs these forms of synaptic enhancement. Using two KI mouse models harboring point mutations that disrupt the function of the Munc13-1 C_2_B and CaM-binding domains, we demonstrate differential effects on the amplitude and time course of STP. DNmut mice − which lack Ca^2+^-phospholipid binding to the Munc13-1 C_2_B domain − showed the most profound STP deficits at hMF synapses. These mice also exhibited significantly impaired working memory formation, supporting the notion that Munc13-1-mediated STP is critical for cognitive function.

**Ca^2+^-dependent upregulation of Munc13-1 function shapes facilitation at hMF synapses** hMF → CA3 pyramidal cell synapses have been described as ’detonator synapses’ because single hMF inputs can reliably trigger postsynaptic CA3 neuron firing^9,34^. Bursts of 2–70 APs induce various forms of presynaptic facilitation − PPF, FF and PTP − at these synapses^3,39^. During presynaptic high-frequency AP firing, presynaptic Ca^2+^ buffer saturation can cause synaptic facilitation because it increases the peak of Ca^2+^ concentrations within local domains triggering glutamate release^5–7^. However, individual AP-induced Ca^2+^ transients at hMF boutons decay within ∼490 ms^30^, so that Ca^2+^ buffer saturation is unlikely to contribute to FF during low-frequency stimulation or to PTP, which lasts several tens of seconds.

An enlargement of the readily releasable pool (RRP) of SVs was previously found to contribute to PTP^9^. We show here that Ca^2+^-dependent regulation of Munc13-1 co-determines several forms of STF in hMF synapses, consistent with the notion that dynamic changes in SV priming contribute to these forms of synaptic facilitation^40,41^. It is conceivable that the number of releases sites occupied with fully primed and fusion-competent SVs is low in resting hMF synapses. During stimulation, this fraction may increase due to the activity-dependent augmentation of SV priming activity, leading to RRP enlargement^40^. Munc13-1 is the prime candidate for mediating such augmentation of SV priming, consistent with the impaired synaptic facilitation in DNmut and WRmut synapses.

While DNmut and WRmut synapses exhibited similarly impaired STF magnitudes across various stimulation frequencies, indicating that the C_2_B and the CaM-binding domain are both essential for facilitation of hMF → CA3 transmission, the buildup of facilitation occurred more slowly in DNmut synapses. Moreover, while LFS-induced facilitation was reduced in WRmut synapses, it was abolished in DNmut synapses. These findings indicate that signaling via the Munc13-1 C_2_B domain is particularly crucial during activity-dependent synaptic enhancement.

### Ca^2+^-dependent upregulation of Munc13-1 activity is required for PTP induction at hMF synapses

An increased RRP contributes to PTP at several types of brain synapses^9–11^ but the underlying signaling pathways and molecular mechanisms are controversially discussed. The data presented here establish that PTP at hMF→CA3 pyramidal cell synapses relies on Ca^2+^-dependent regulation of Munc13-1. Genetic manipulations of the two regulatory pathways converging on Munc13-1 differentially affect PTP induction threshold and magnitude. Thus, Ca^2+^-phospholipid and Ca^2+^-CaM signaling through Munc13-1 may be differentially recruited during different presynaptic activity patterns. Ca^2+^-dependent regulation of Munc13-1 likely acts in cooperation with other signaling pathways, such as DAG binding to the Munc13-1 C_1_ domain, or in addition to other priming proteins, such as Munc13-2 − whose expression varies across hMF synapses (**Figure S3B**). Variability in Munc13-1 and Munc13-2 expression levels may underlie the heterogeneity of PTP among wt hMF synapses and contribute to differences in PTP impairment observed in individual DNmut and WRmut synapses.

### Linking Munc13-1-dependent short-term plasticity to working memory

Alongside the STF and PTP deficits in hMF → CA3 pyramidal cell synapses, DNmut mice exhibited pronounced deficits in the 8ARM working memory task. In particular, these mice were severely impaired in recalling prior reward retrievals, often re-entering rewarded food zones repeatedly and spending five times longer there than wt mice. Although these behavioral tests cannot definitively resolve the cellular mechanisms involved, our data indicate that disrupted STP may impair the transient memory traces needed for short-term decision-making. These findings support recent models proposing that synaptic alterations, in addition to persistent neuronal firing, contribute to working memory storage^2,36,42^.

While WRmut and DNmut hMF synapses displayed similar STF deficits during train stimulation and similarly impaired PTP in response to longer induction trains, DNmut hMF synapses additionally required stronger stimulation for PTP induction. When assayed by the 8ARM task, WRmut mice performed comparably to wt early in training but showed a slight increase in time to task completion during later sessions, hinting at impairments in reward retrieval processing similar to those in DNmut mice. These findings indicate that Ca^2+^-CaM and Ca^2+^-phospholipid signaling to Munc13-1 are both essential for working memory-relevant synaptic plasticity. The milder behavioral phenotype of WR mutants as compared to the DN mutants is likely related to the milder synaptic plasticity defects.

Future studies employing conditional KI mice will help to further strengthen the connection between specific synapse types and the observed behavioral phenotypes. Although we cannot unequivocally establish an association between the impaired function of hMF → CA3 pyramidal cell synapses and the behavioral deficits, our data provide the first experimental evidence of a link between Munc13-1-regulated SV priming, STP, and working memory.

## Conclusion

Our results demonstrate that the Ca^2+^-dependent regulation of Munc13-1 via the C_2_B and the CaM-binding domains is not only essential for STP and PTP at hMF synapses, but also plays a key role in supporting working memory formation. Together, these findings establish a mechanistic link between presynaptic modulation of glutamate release and higher-order cognitive processes, highlighting the importance of Munc13-1-mediated SV priming and temporary storage of information.

## METHODS

### Mouse lines

The Munc13-1 knock-in (KI) mice were generated as reported^16,17^ and maintained and bred in the animal facility of the Max Planck Institute of Multidisciplinary Sciences. For electrophysiological experiments, mouse lines were maintained on a C57BL/6N background, and experimental groups consisting of male or female WT and KI littermates were generated from heterozygous breeding pairs. For behavioral experiments, all mouse lines were backcrossed for six generations onto a C57BL/6J background to avoid the retinal dysplasia phenotype observed in the C57BL/6N strain. For each mouse line, experimental groups consisting of male WT and KI littermates were generated from heterozygous breeding pairs, and behavioral testing was initiated at the age of 8–12 weeks. Animals were maintained on a 12 hours-light/dark cycle (7 am / 7 pm), with food and water ad libitum (except during the 8-arm radial maze test as described below), and all experiments were performed during the light cycle. All procedures were approved by the State of Lower Saxony (Landesamt für Verbraucherschutz und Lebensmittelsicherheit, TVA 33.19-42502-04-21/3798) and were carried out in agreement with the guidelines for the welfare of experimental animals issued by the Federal Government of Germany and the Max Planck Society.

### Immunostaining

Immunostaining experiments were performed on P21-25 horizontal sections containing the hippocampal mossy fiber axon track, using antibodies against Munc13-1 (rabbit polyclonal, Synaptic Systems #126102), Munc13-2 (rabbit polyclonal, Synaptic Systems #126203), vGlut1 (guinea pig polyclonal, Synaptic Systems #135304), and synaptoporin (rabbit polyclonal, Synaptic Systems #102002). Brains perfused with 4% PFA and immersed in 30% sucrose solution in 0.1 M PB were cut into 15-µm-thick horizontal cryosections of the hippocampal region and transferred to 0.1 M PB solution. Floating sections were incubated for 90 min at RT in blocking solution (0.1 M PB, 10% normal goat serum, 0.5% Triton X-100, pH 7.4) before being treated overnight at 4°C with the primary antibodies diluted in incubation buffer (0.1 M PB, 5% normal goat serum, 0.3% Triton X-100, pH 7.4). After washing in PB, sections were incubated for 2 h at RT in the dark with fluorescent secondary antibodies (Alexa 488-coupled goat anti-rabbit, Alexa-555-coupled goat anti-guinea pig, Alexa-647-coupled goat anti-guinea pig, 1:500; Invitrogen) and DAPI, diluted in incubation buffer. Coverslips were mounted with Aqua-PolyMount (Polysciences). For immunolabeling experiments, in which the antibody against Munc13-2 was employed, TBS-1x buffer was used instead of PB. Confocal laser scanning micrographs of presynaptic compartments at the *stratum lucidum* of the hippocampus were acquired with a Carl Zeiss LSM880 confocal microscope. An 40X oil-immersion objective (NA = 1.3) and a pinhole setting of 1 AU were used to obtain single-plane micrographs (1024 pixels × 1024 pixels; x-y pixel spacing of 0.29 nm) in sequential scanning mode. Laser power and gain were adjusted to ensure that signals were in the linear range of detection. We performed line profile analysis of Munc13-1 and Munc13-2 fluorescence intensities of vGlut1-positive puncta within the *stratum lucidum*.

### Slice preparation

Acute horizontal slices of postnatal (P18-30) mice of either sex were used for electrophysiology. After decapitation, brains were dissected and immersed in ice-cold low-Ca^2+^ and high-sucrose artificial CSF (aCSF) containing (in mM): 87 NaCl, 2.5 KCl, 25 NaHCO_3_, 1.25 NaH_2_PO_4_, 25 glucose, 0.5 CaCl_2_, 7 MgCl_2_, 75 sucrose (pH 7.4, bubbled with 95% O_2_, 5% CO_2_). Brains were glued onto the stage of a VT1000S vibratome (Leica), and 290 µm-thick horizontal slices containing the CA3 region of the hippocampus were cut. Slices were incubated for 40 min at 35°C in a chamber containing the same solution as the one used for dissection and slicing. Slices were kept in this solution at RT (21–24°C) for up to 5 h after recovery.

### Electrophysiology

Whole-cell patch-clamp recordings were made from pyramidal neurons of the CA3 region of the hippocampus at near physiological temperature using an EPC-10 amplifier controlled by Pulse or PatchMaster software (HEKA Elektronik). Sampling intervals and filter settings were 20 μs and 5 kHz, respectively. For voltage-clamp recordings, the membrane potential was set to −70 mV. Slices were placed into the recording chamber and allowed to wash for 10 min in normal aCSF before recordings commenced. Normal aCSF contained (in mM): 125 NaCl, 2.5 KCl, 25 NaHCO_3_, 1.25 NaH_2_PO_4_, 25 glucose, 2 CaCl_2_, 1 MgCl_2_ (pH 7.4, bubbled with 95% O_2_, 5% CO_2_). Neurons were visualized using near IR illumination through a 40× (Zeiss Achroplan, 0.8 NA Water immersion) objective. Perfused solution was warmed by a Single Inline Solution Heater SH-27B, controlled by a Single-Channel Temperature Controller TC324C (Multichannel Systems).

Patch pipettes were pulled from borosilicate glass capillaries with filament (Science Products) to have an open-tip resistance of 3–5 MΩ when filled with intracellular solution. For voltage-clamp recordings the intracellular solution contained (in mM): 120 K-gluconate, 30 KCl, 10 EGTA, 2 MgCl_2_, 2 Na-ATP, 10 HEPES, 2 QX-314, 314 mOsm, pH 7.3. Some cells were filled with intracellular solution containing 0.4% biocytin for light microscopic characterization of the morphology of CA3 pyramidal cells in the slice (**Figure S1**). We did not correct for the liquid junction potential.

During voltage-clamp recordings, aCSF was supplemented with 25 μM bicuculline methiodide (HelloBio) to block GABA_A_R-mediated IPSCs. Mossy fibers were stimulated with a glass electrode filled with aCSF and placed in stratum lucidum ≥80 μm away from the cell body. Transmitter release was evoked by applying brief electrical pulses (100 μs, 10–50 V). Stimulus intensity was adjusted to obtain peak EPSC amplitudes corresponding to single-axon stimulation, assuming that mean putative unitary EPSC amplitudes have a range of 40–500 pA, and exhibited facilitation during stimulus trains (∼5-fold after ∼10 APs; **Figures S1C**). eEPSCs were blocked by 2 µM DCG-IV and by 20 µM of NBQX (**Figures S1B and S1C**), as expected for hMF inputs.

All offline analysis of electrophysiological data was performed in Igor Pro (Wavemetrics). Voltage-clamp recordings were leak-corrected and low-pass filtered with a cut-off frequency fc=3 kHz using a 10-pole digital Bessel filter.

### Behavioral testing

Each cohort of mice (male WT and KI littermates, 8–12 weeks old at the beginning of testing) was assessed on three behavioral paradigms, a visual cliff (VC) test, an open field (OF) test and an 8-arm radial maze (8ARM) test. VC and OF testing were conducted in week 1 of the behavioral battery, with an interval of at least 48 hours between the tests, while 8ARM testing was conducted in weeks 2 and 3. Behavioral testing chambers were custom-built by the precision mechanics machine shop of the Max Planck Institute of Multidisciplinary Sciences. Performance of the animals in the testing chambers was recorded using an overhead camera system and scored automatically using the Viewer software (Biobserve, St. Augustin, Germany). Experimenters were blind to genotype at all stages of data acquisition and analysis. Between each mouse, the arena was cleaned thoroughly with 70% ethanol followed by water to eliminate any odors left by the previous mouse.

*Visual cliff test*. The VC test was conducted in a rectangular arena made of clear acrylic glass (40 x 60 cm surface area with 15 cm high walls, light intensity 20 lux), which was placed on a metal frame at a height of 50 cm above a black-and-white patterned floor. One half of the arena was underlaid with an opaque base consisting of the same black-and-white patterned material (‘platform’, 40 cm × 30 cm), while other half was left clear to create the optical illusion of a visual cliff (‘cliff’, 40 cm × 30 cm). Mice were placed in the center of the opaque platform and were permitted to explore the arena for 3 min. Parameters assessed included time in the platform and cliff zones.

*Open field test*. The OF test was conducted in a square arena made of white plastic (50 cm × 50 cm surface area with 50 cm high walls, light intensity 30 lux). Mice were placed in the corner of the OF and were permitted to explore the arena for 10 min. Three zones were defined in the Biobserve software (see also **Figure S4B**), a center zone (25 cm × 25 cm), an intermediate zone (37.5 cm × 37.5 cm excluding the center), and a peripheral zone (the remaining outermost zone of the arena). Parameters assessed by the software included the total distance traveled across all zones and the time spent in each zone.

*8-arm radial maze*. The 8ARM test was conducted in an apparatus consisting of a central circular chamber (diameter 22 cm) with eight protruding arms (35 cm × 6 cm with 15 cm high walls, light intensity 40 lux). The floor of the apparatus was made of grey plastic, while the walls were made of clear Polymethyl methacrylate to facilitate spatial orientation. The end of each arm contained a food zone (6 cm × 6 cm as defined by the Biobserve software), into which a food receptacle with a shallow circular indentation (diameter 1.6 cm, depth 0.5 cm) was placed. High-contrast visual cues were attached to the end wall of each arm above the food receptacle, with distinct patterns for each arm. Additional visual cues were attached to the walls of the behavioral testing room at a distance of approximately 1–2 m from the 8ARM apparatus. Behavioral testing was conducted across 10 sessions (5 daily sessions per week for two weeks, separated by the weekend). Throughout this period, mice were maintained on a food restriction schedule, in which they received free access to food for 1.5 hours per day following behavioral testing, as well as ad libitum food over the weekend (Friday afternoon to Sunday afternoon). Mice were weighed 3× per week to ensure maintenance of >85% free-feeding body weight. Prior to the start of behavioral testing, mice were additionally habituated for three days to the food rewards used in the behavioral paradigm (‘Dustless Precision Pellets’, Bio-Serv, sucrose 20 mg). Each daily behavioral session consisted of four trials lasting up to 4 min, with an inter-trial interval of 1 hour. Prior to each trial, three of the eight food zones were baited with a food reward. The location of the rewarded food zones was kept constant for each mouse across all sessions and trials, but was counterbalanced between mice to control for spatial bias. To initiate each trial, mice were placed into the central chamber and allowed to freely explore the apparatus. Trials were terminated when the mouse had retrieved all three food rewards, or after 4 min if the food rewards were not all retrieved. The following parameters were assessed (averaged across the four daily trials): (1) Number of working memory errors, defined as repeated entries into a previously rewarded food zone after the reward had already been retrieved, (2) number of reference memory errors, defined as entries into a non-rewarded food zone, (3) average time spent in rewarded food zone, and (4) total distance travelled.

### Statistical analysis

*Functional data.* Data are expressed as mean ± standard error of the mean (SEM). Error bars in all graphs indicate SEM. In figure legends, ’n’ refers to the number of CA3 cells recorded (**Figures 1-3**, **Figures S1-S3**). Because the plots of samples distributed following a Gaussian distribution, statistical significance of differences in sample location was tested with one-way ANOVA test followed by Tukey test (**Figures 1-3**), or paired Student’s t-test (**Figure S3**). Statistical analysis was conducted using Igor Pro and R (R Foundation for Statistical Computing). One, two, and three asterisks in figure panels indicate statistically significant differences at p< 0.05, p< 0.01, and p< 0.001, respectively.

*Behavioral data.* Statistical analysis was conducted using Prism (GraphPad Software, La Jolla, CA, USA). Outliers were identified using the Grubbs test (alpha = 0.01). Data to assess time in zone in the visual cliff (**Figures S4D-E**) and open field test (**Figures S4H-I**) were subjected to two-way repeated-measures ANOVA, with zone as the repeated factor and genotype as the second factor. Data to assess working memory errors (**Figures 4D-E**), total distance travelled (**Figures 4G-H**), time to trial completion (**Figures 4J-K**) and time in reward zone (**Figures 4M-N**) in the 8-arm radial maze were subjected to two-way repeated-measures ANOVA, with training day as the repeated factor and genotype as the second factor. Significant main effects of zone/day or genotype, as well as significant zone/day x genotype interactions, are indicated in the title of each graph as described in the corresponding figure legend (except where stated otherwise). Post-hoc analysis was conducted using Tukeýs test for comparison between groups, and significant differences are indicated with asterisks above the corresponding comparison. Data to assess distance travelled in the open field (**Figures S4F-G**), as well as comparisons of the last three days of testing across different lines (**Figures 4F, 4I, 4L, 4O**) were subjected to an unpaired two-tailed Student’s t-test (if normally distributed as assessed by the Shapiro-Wilk test) or a Mann-Whitney test (if not normally distributed). Significant effects are indicated with asterisks above the corresponding bars.

## Author contributions

F.J.L.-M. designed and performed electrophysiological experiments with the assistance of H.T.; F.J.L.-M. and H.T. analyzed electrophysiological data; H.T. provided software routines for analysis; F.J.L.-M. and T.L.-H. performed immunofluorescence experiments; D.K.B. designed and supervised behavioral experiments; S.W. conducted behavioral tests with the assistance of D.K.B.; D.K.B. analyzed behavioral data; N.L. generated and validated Munc13-1 KI mouse lines; F.J.L.-M., D.K.B., N.L., H.T. and N.B. wrote the paper.

## Acknowledgements

We thank the MPI-NAT Animal Facility for mouse husbandry, the MPI-NAT AGCT laboratory for genotyping, and Markus Krohn and the MPI-NAT Precision Mechanics Workshop for the design and construction of the behavioral testing chambers. This work was supported by an Alexander von Humboldt Foundation postdoctoral fellowship (F.J.L.-M.), by the Agencia Estatal de Investigación (PID2022-141685NA-I00, F.J.L.-M.; PID2021-128845NA-I00, T.L.-H.), by the Max Planck Partner Group Program (F.J.L.-M.), and by the German Research Foundation (Heisenberg Grant KR 5239/1-1, D.K.B.; SFB 1286/A9, N.B. and 1286/A11, N.L.; Br1107/15, N.B.; EXC-2049-3906880787, N.L.) and by the Target ALS Foundation (N.L.).

## SUPPLEMENTAL FIGURE LEGENDS

**Supplemental Figure 1.**
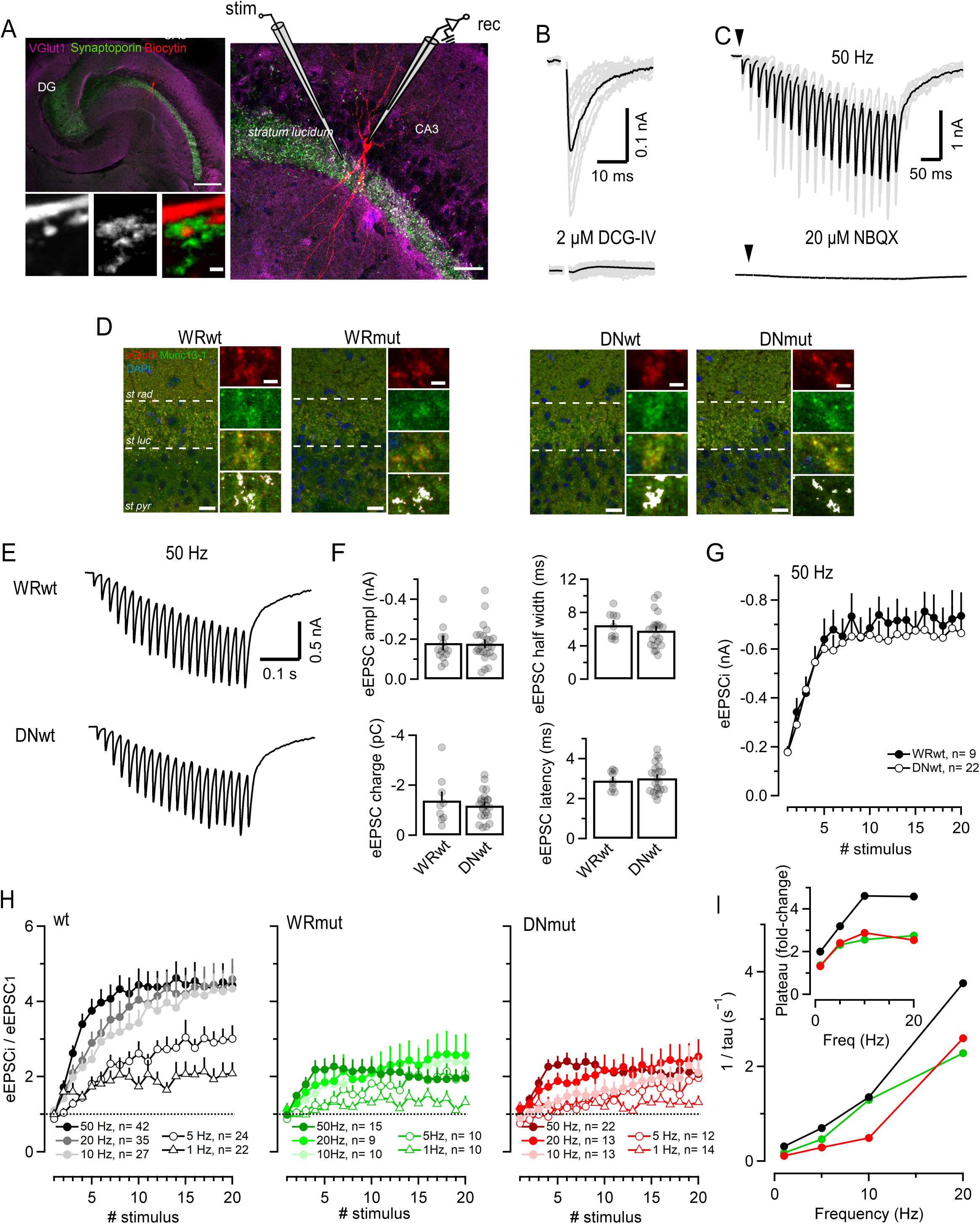
Identification of hMF eEPSCs, Munc13 expression in hMF terminals and short-term plasticity in wt hMF synapses (Related to Figure 1). **(A)** Confocal fluorescence microscopy images and experimental set up for electrophysiology. A CA3 pyramidal neuron was recorded in whole-cell configuration using a glass pipette containing the recording electrode (*rec*). The stimulation electrode (*stim*) was positioned ∼100 µm away from the soma of the patched CA3 neuron in the *stratum lucidum*. Electrical stimuli (100 µs duration) with intensities above the threshold of action potential (AP) generation of mossy fiber (hMF) axons elicited putative unitary excitatory postsynaptic currents (eEPSC) in CA3 pyramidal neurons. eEPSC were pharmacologically isolated by the presence of 25 μM bicuculline methiodide in the bath to block IPSCs. Confocal images show a hippocampal slice used for electrophysiology, subsequently fixed and immunolabeled against the presynaptic vesicular transporter of glutamate type 1 (vGlut1, *magenta*) and synaptoporin (*green*), a specific marker of hMF boutons. During recording, a CA3 pyramidal neuron was filled with 0.4% biocytin via the recording pipette. Streptavidin conjugated to Alexa-Fluor 555 (*red*) was included in the secondary antibody cocktail for biocytin labelling during immunostaining. The low-magnification image (10x, *top left*) displays the hippocampus, containing the DG and CA3 areas (scale bar, 200 µm). A higher magnification image (40x, *right*) highlights a patched CA3 pyramidal cell filled with biocytin and the experimental setup (scale bar, 40 µm). The high-magnification inset (*bottom left*) shows a postsynaptic spine (biocytin, *red*) surrounded by a synaptoporin-positive presynaptic structure (*green*), presumably corresponding to an hMF synapse (scale bar = 1 µm). **(B)** Unitary eEPSCs recorded in a CA3 pyramidal (as in panel A) in response to electrical stimulation of a putative single hMF axon. Top: representative traces (12 repetitions at 0.5 Hz; *light traces*) with the average eEPSC superimposed (*solid trace*). Bottom: selective blocking of hMF eEPSCs with 2 µM DCG-IV (*bottom*). **(C)** Short-term facilitation of eEPSC trains in response to 20 stimuli at a 50 Hz. Top: nine repetitions (*light traces*) superimposed on the average eEPSC train (*solid trace*). Bottom: selective blocking of eEPSCs by 20 µM of NBQX, the selective and competitive AMPA/kainate receptor antagonist. Arrowheads indicate the onset of the stimulus train. **(D)** Confocal images showing Munc13-1 (*green*), vGlut1 (*red*), and DAPI (*blue*) in the *stratum lucidum* of hippocampal sections from P23 WRwt and WRmut mice (*left*), and P21 DNwt and DNmut mice (*right*). The CA3 sub-regions are indicated: *stratum radiatum* (*st rad*), *stratum lucidum* (*st luc*) and stratum pyramidale (*st pyr*) (scale bar 20 µm). Insets (*right*): magnified views of vGlut-1 (*top*), Munc13-1 (*second*), merged signals (*third*), and areas of co-localization (*white*) at putative hMF boutons (*bottom*) (scale bar = 5 µm). **(E)** Average eEPSC trains recorded in wt synapses of the two KI lines (WRwt, n=8; DNwt, n=19) in response to 20 stimuli at 50 Hz. **(F)** Bar graphs and scatter dot showing eEPSC parameters recorded in wt synapses of the two KI lines: amplitude (*top left*), charge (*bottom left*), half-width (*top right*) and latency (*bottom right*). **(G)** Summary of eEPSC amplitudes recorded during 20-stimulus trains at 50 Hz in WRwt (*solid circles*) and DNwt (*open circles*) hMF synapses. **(H)** Average normalized eEPSC amplitudes recorded in wt (*black*, pooled from wt synapses of the two KI lines), WRmut (*green*) and DNmut (*red*) synapses in response to 20-stimulus trains at varying frequencies (1, 5, 10, 20 and 50 Hz). Normalization was performed by dividing each eEPSC by the mean eEPSC_1_ recorded at any frequency in the same synapse. Dotted lines indicate unity. **(I)** Representation of the inverse of the time constant during enhancement of facilitation as a function of the stimulation frequency (1, 5, 10, 20 Hz) for wt, WRmut and DNmut hMF synapses. The time constant τ was determined from a fit to the function y = 1 + (A−1) · (1−exp(−t/τ)) where y represents the normalized eEPSC amplitude as shown in (H) and t is the time since stimulation onset. The inset shows the change in plateau (value of A in the fit function) during facilitation as a function of stimulation frequency. 50 Hz eEPSC trains were excluded from the analysis because facilitation time courses could not be fitted with a single exponential function in WRmut and DNmut synapses, unlike in wt hMF synapses. Error bars indicate SEM.

**Supplemental Figure 2.**
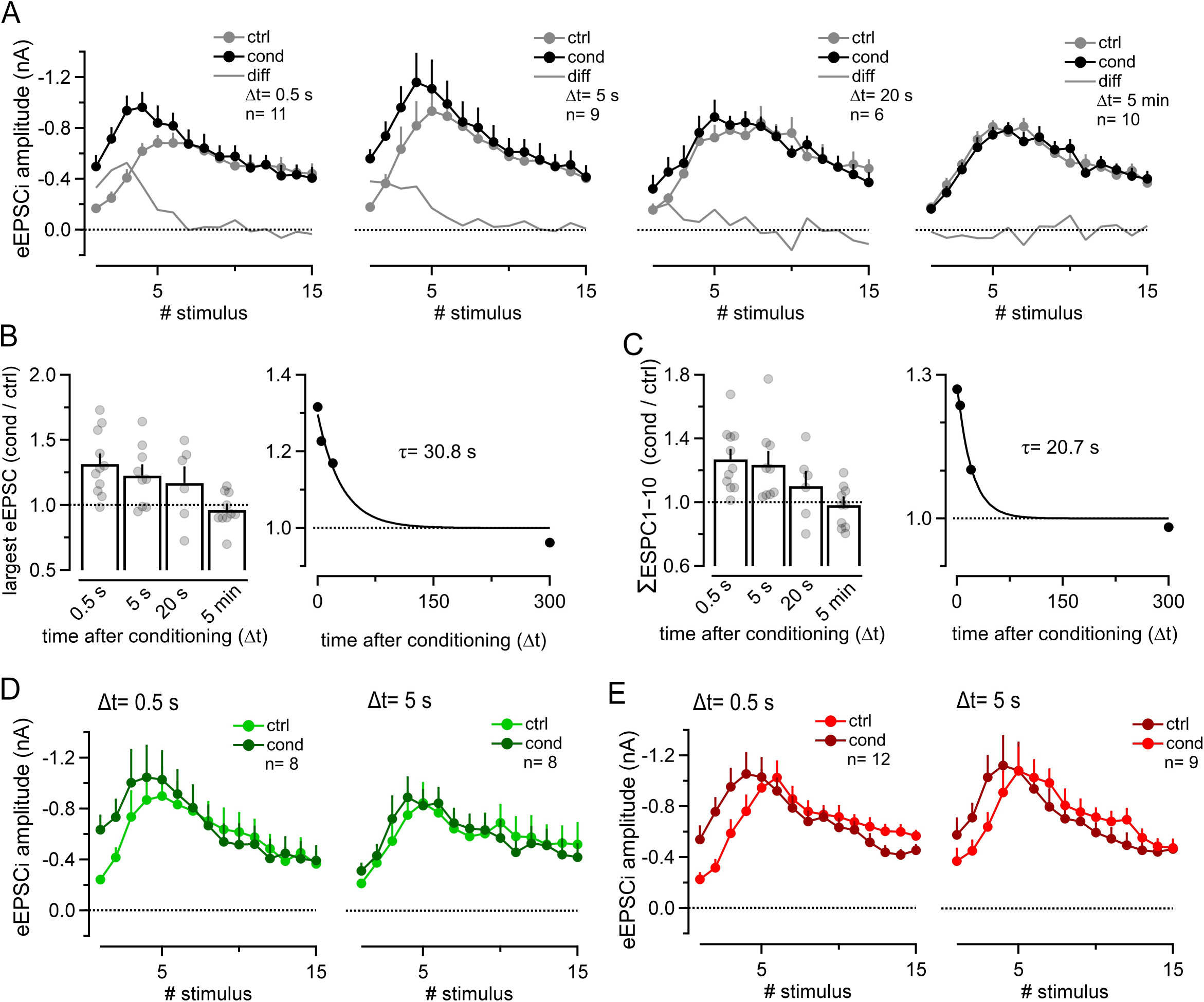
Neurotransmitter release transiently potentiates in hMF synapses following low-frequency stimulation pre-conditioning (Related to Figure 2). **(A)** Summary data of eEPSC amplitudes recorded in wt hMF synapses in response to 15-stimulus trains at 100 Hz, delivered before (ctrl, *gray*) and after (cond, *black*) LFS. Each plot represents average data from synapses where the cond train was recorded at different intervals (Δt) after LFS: 0.5 s (*left*), 5 s (*middle left*), 20 s (*middle right*) and 5 min (*right*). eEPSC differences (eEPSC *diff*) between the corresponding cond and ctrl trains in wt synapses in which cond trains were recorded at varying Δt (0.5 s, 5 s, 20 s, 5 min) after LFS are plotted. eEPSC *diff* was calculated by substracting the mean eEPSC amplitude for each stimulus between the cond and ctrl trains. Dotted lines indicate zero difference. **(B)** Bar graphs and scatter dot plots showing the ratio of the largest eEPSC amplitudes between the cond and ctrl trains as a function of the interval (Δt) after LFS (*left*). Dotted lines represent unity. The average values were plotted against time after LFS and fitted to an exponential decay function with a time constant of ∼31 s (*right*). **(C)** Bar graphs and scatter dot plots showing the ratio of cumulative eEPSC amplitudes (first 10 responses of the train) between the cond and ctrl trains, as a function of Δt after LFS (*left*). The average values were plotted against time after LFS and fitted to an exponential decay function with a time constant of ∼21 s. Time constant values from (C) and (D) could be interpreted as the de-priming time constant in wt hMF synapses which ranges 20–30 ms. Dotted lines represent unity. **(E-F)** Summary data of eEPSC amplitudes recorded in WRmut (E) and DNmut (F) hMF synapses in response to 15-stimulus trains at 100 Hz, delivered before (ctrl, *light color*) and after (cond, *solid color*) LFS. Each plot represents average data from synapses where the cond train was recorded at different intervals (Δt) after LFS: 0.5 s (*left*) and 5 s (*right*). Dotted lines indicate zero difference.

**Supplemental Figure 3.**
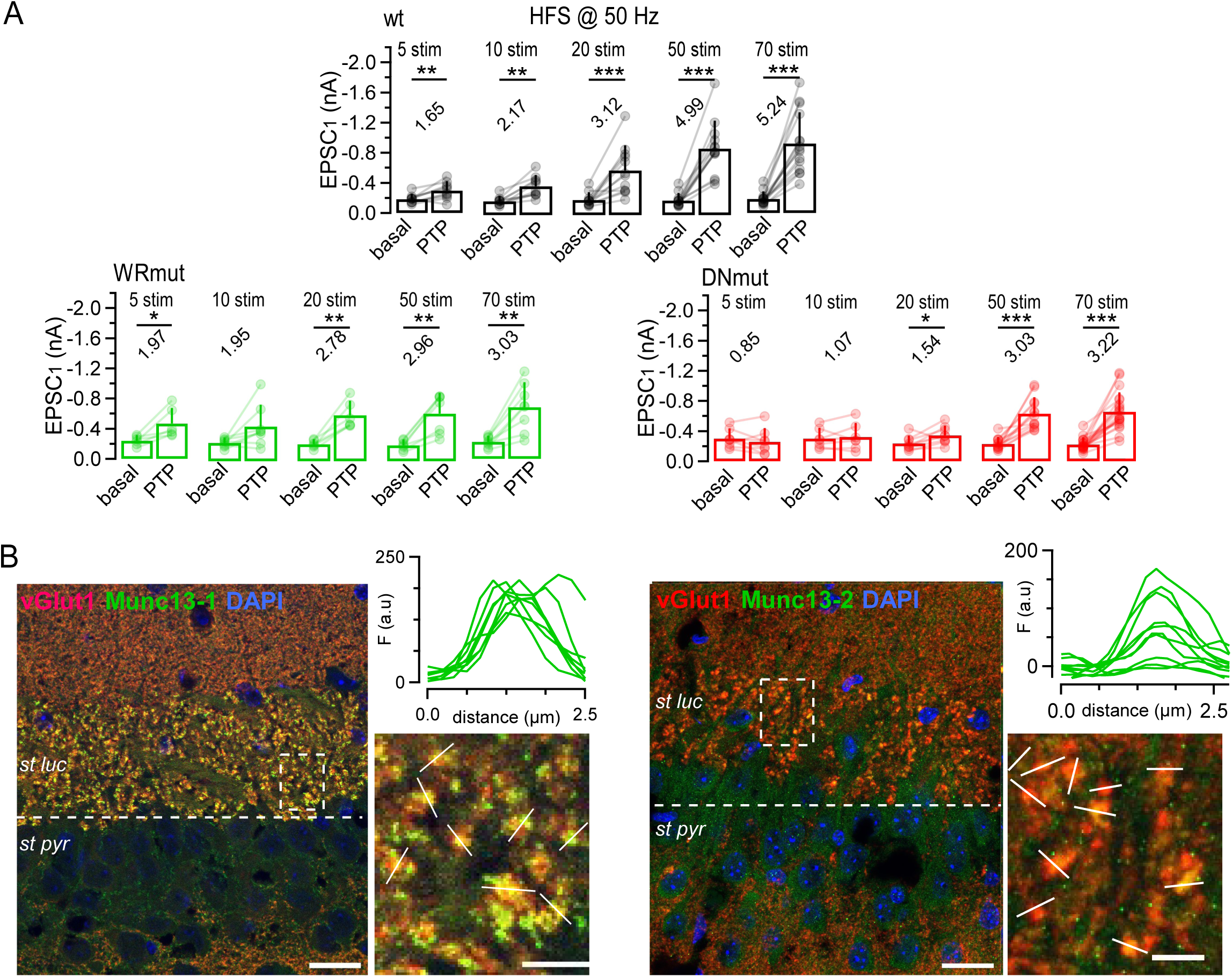
Required cooperation between C_2_B and CaM binding domains of Munc13-1 to potentiate SV release after synapse conditioning with high-frequency stimulation (Related to Figure 3). **(A)** Summary bar graph and scatter dot plots comparing the effects of HFS with varying number of stimuli on unitary mossy fiber eEPSCs. For each cell, the eEPSC_1_ amplitude in response to PP@20ms is compared before (basal) and after (PTP) HFS. Data are shown for wt (*black, top*), WRmut (*green, bottom left*), and DN mut (*red, bottom right*) hMF synapses. **(B)** Presynaptic anti-Munc13-1 and anti-Munc13-2 fluorescence in hMF boutons of hippocampal sections obtained from wt mice. Maximum-intensity projections were obtained from stacks of 2–3 confocal images of P25 hippocampi. Presynaptic compartments were identified by anti-vGluT1 immunolabeling (*red colour in left column, magenta colour in the right column*). Sections were co-stained with either anti-Munc13-1 (*green colour in the left column*) or anti-Munc13-2 antibodies (*yellow colour in right column*) to demonstrate presence or absence of these isoforms in presynaptic glutamatergic synapses of wt mice. DAPI stains nuclei. The CA3 sub-regions are indicated: *stratum lucidum* (*st luc*) and stratum pyramidale (*st pyr*) (scale bar = 20 µm). Insets (*right*) magnify views of merged signals at putative hMF boutons of the stratum lucidum (scale bar = 5 µm). Note that anti-Munc13-1 immunofluorescence is found in virtually all putative hMF synapses, while anti-Munc13-2 immunofluorescence is absent from some putative hMF synapses . Error bars indicate SEM. (Paired t-test, * p<0.05, ** p<0.01, *** p<0.001).

**Supplemental Figure 4.**
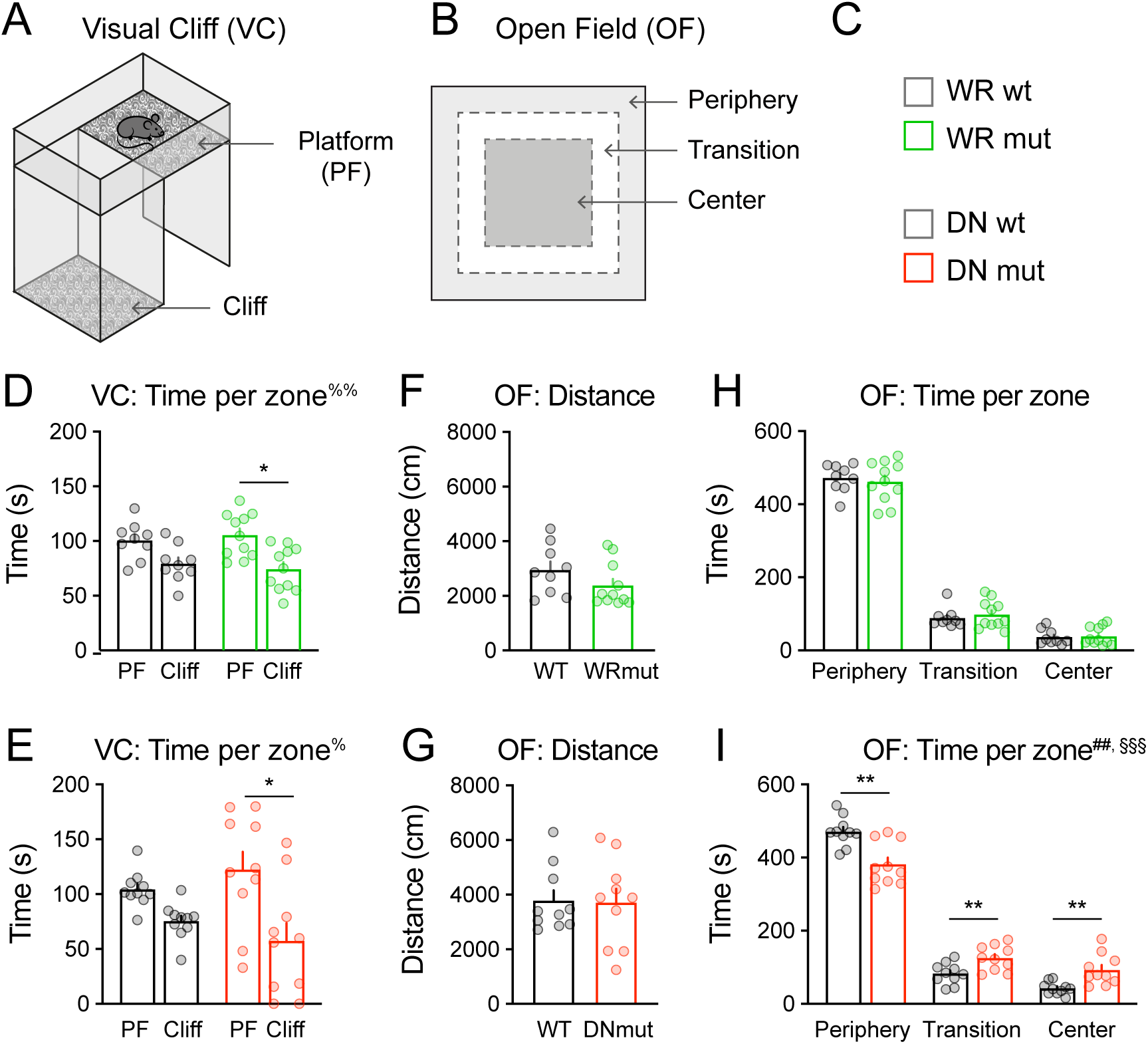
Normal visual performance of WRmut and DNmut mice (Related to Figure 4). **(A-B)** Schematic representations of the visual cliff paradigm (A) and the open field paradigm (B). **(C)** Representation of behavioral cohorts tested. **(D-E)** Time spent in platform (PF) vs. cliff zones in the VC paradigm for WR (D) and DN (E) mouse lines. A difference in time spent in the platform vs. cliff zones indicates that mice were able to visually distinguish between these zones. Two-way ANOVA with repeated measures for zone (main effect of zone, ^%^ p<0.05, ^%%^ p<0.01; main effect of genotype, not significant; genotype x zone interaction, ^§^ p<0.05) followed by Tukey’s *posthoc* test (* p<0.05). (F-G) Total distance travelled during the OF test for WR (F) and DN (G) mouse lines. Unpaired two-tailed Student’s t-test or Mann-Whitney test. (H-I) Time spent in the periphery, transition and center zones of the OF paradigm for WR (H) and DN (I) mouse lines. Two-way ANOVA with repeated measures for zone (main effect of zone, not shown; main effect of genotype, ^##^ p<0.01; genotype x zone interaction, ^§§§^ p<0.001) followed by Tukey’s *posthoc* test (** p<0.01). Error bars represent SEM.

